# CRISPR-Cas9 targeted deletion of the *C9orf72* repeat expansion mutation corrects cellular phenotypes in patient-derived iPS cells

**DOI:** 10.1101/051193

**Authors:** Mochtar Pribadi, Zhongan Yang, Tanya S. Kim, Elliot W. Swartz, Alden Y. Huang, Jason A. Chen, Deepika Dokuru, Jaeyun Baek, Fuying Gao, Andrea T. Fua, Kevin Wojta, Qing Wang, Anna Karydas, Jamie Fong, Ed Lezcano, Stephanie Ng, Farid F. Chehab, Harry V. Vinters, Bruce L. Miller, Giovanni Coppola

**Affiliations:** Departments of Psychiatry, David Geffen School of Medicine, University of California at Los Angeles, Los Angeles, CA, USA; Memory and Aging Center, University of California San Francisco, San Francisco, CA, USA; Department of Laboratory Medicine, University of California San Francisco, San Francisco, CA, USA.; Neurology, Semel Institute for Neuroscience and Human Behavior, David Geffen School of Medicine, University of California at Los Angeles, Los Angeles, CA, USA; Department of Pathology and Laboratory Medicine, David Geffen School of Medicine, University of California at Los Angeles, Los Angeles, CA, USA.

## Abstract

The large hexanucleotide (GGGGCC) repeat expansion in the non-coding promoter region of *C9orf72* is the leading cause of Frontotemporal Dementia (FTD) and Amyotrophic Lateral Sclerosis (ALS). Mechanisms underlying neurodegeneration are not clear, and both a C9orf72 loss of function and a gain of toxicity, in the form of RNA foci or dipeptide repeat deposition, are implicated. CRISPR (clustered regularly interspaced short palindromic repeats)-Cas9-mediated genome editing is an attractive strategy for disease modeling and therapeutic intervention. Here we show that this system can be utilized to completely remove the large repeat expansion mutation within *C9orf72* in patient-derived induced pluripotent stem (iPS) cells. Removal of the mutation prevented RNA foci formation and promoter hypermethylation, two phenotypes of the *C9orf72* mutation. Interestingly, these changes did not significantly alter C9orf72 expression at the mRNA or protein level. This work provides a proof-of-principle for the use of CRISPR-Cas9-mediated excision of the pathogenic *C9orf72* repeat expansion as a therapeutic strategy in FTD/ALS.

**One Sentence Summary:** CRISPR-Cas9-mediated excision of the large *C9orf72* repeat expansion mutation rescues RNA foci formation and promoter hypermethylation without altering C9orf72 transcript and protein expression.

## Introduction

The most frequent genetic cause of Frontotemporal Dementia (FTD) and Amyotrophic Lateral Sclerosis (ALS) is a hexanucleotide (GGGGCC) repeat expansion in the chromosome 9 open reading frame 72 (*C9orf72*) gene (*1*, *2*). The repeat lies between two alternative non-coding first exons of this gene known only to have structural similarities to the DENN (differentially expressed in normal and neoplastic cells) family of Rab GEFs (*3*). Affected carriers can have up to hundreds to thousands of repeats, though the number is highly variable, even within an individual (*4*). Normal individuals are generally thought to have less than about 30 repeats, however a definitive threshold for pathogenicity is not currently established (*1*, *2*, *5*).

The precise mechanisms underlying *C9orf72*-linked neurodegeneration remain unclear. The location of the mutation within a non-coding regulatory region points towards haploinsufficiency as a pathogenic mechanism. Early reports indeed showed a reduction in *C9orf72* transcription (*1*, *6*), which was observed in a number of subsequent studies along with decreased protein expression (*7*–*11*) and hypermethylation of both CpG islands and histones proximal to the repeat expansion (*5*, *12*–*14*). Additionally, knockout of the *C9orf72* ortholog in zebrafish and *C. elegans* resulted in motor deficits and neurodegeneration (*11*, *15*). However, deletion of *C9orf72* in mice did not produce neurodegeneration, motor deficits, or decreased survival (*16*), and knockdown of *C9orf72* in human iPS cell-derived neurons did not alter viability (*17*). Together with the lack of any disease-causing coding mutations in the gene, the role of the C9orf72 protein in disease pathogenesis, if any, is still uncertain (*18*).

Transcription of the repeat itself yields an RNA species with extensive secondary structures that aggregate to form nuclear RNA foci (*1*). These foci are thought to confer a toxic gain of function by sequestering essential RNA binding proteins, similar to the sequestration of Muscleblind in myotonic dystrophy (*9*, *17*, *19*–*21*). Furthermore, there is evidence that these repeat-containing RNAs form G-quadruplex structures which disrupt transcription and nucleocytoplasmic transport (*22*, *23*). However, a *Drosophila* model expressing exclusively *C9orf72* RNA foci showed very little toxicity (*24*).

Repeat-associated non-canonical (RAN) translation, where the repeat is translated in all three reading frames independent of a start site, is now known to occur in most repeat expansion disorders (*25*). RAN translation of the hexanucleotide repeat in *C9orf72* produces poly-dipeptide chains of GP, GA, and GR from the sense transcript and PA, PR, and PG from the antisense strand (*26*–*28*). In the diseased brain, these dipeptides form p62-positive intracellular inclusions reminiscent of other neurodegenerative proteinopathies, however studies vary on their toxicity and relevance to pathogenesis. Overexpression in cell cultures revealed a greater aggregation potential for the GA species, inducing neuronal toxicity through ER stress, and disruption of the proteasomal pathway (*29*, *30*). In *Drosophila*, GA expression is relatively innocuous while the arginine-containing species, GR and PR, induced striking neurodegeneration (*31*–*33*). Interestingly, human patient expression of the dipeptide products does not seem to correlate well with the pattern of degeneration or clinical phenotype (*34*–*36*). In fact, dipeptide deposition and foci formation seem to appear long before clinical symptoms manifest, and are also present in clinically normal patients who have only about 30 repeats (*37*, *38*). Furthermore, the recent creation of BAC transgenic mice by two independent labs showed the presence of the foci and dipeptide inclusions without neurodegeneration or any behavioral or motor phenotypes (*39*, *40*).

Regardless of the mechanisms, it remains clear that the repeat expansion in *C9orf72* is responsible for the majority of FTD and ALS, up to 6% of sporadic cases and 25% and 40% of familial cases of FTD and ALS respectively *(41)*. Advances in genome editing with the availability of CRISPR technology bring wide possibilities in disease modeling and gene therapy by enabling precise and efficient DNA double-stranded breaks *(42)*. Recently, this system was shown in vivo to partially rescue deficits in a mouse model of Duchenne muscular dystrophy *(43*–*45*). Here we show the excision of the full hexanucleotide repeat expansion from FTD patient-derived iPS cells using the CRISPR-Cas9 system. Deletion of the expanded repeat rescued RNA foci formation and the promoter hypermethylation observed in repeat expansion carrier cells, while having little effect on the transcription and protein expression of *C9orf72*.

## Results

### Generation and characterization of iPS cell lines

Human dermal fibroblasts from *C9orf72* repeat-positive FTD/ALS patients were reprogrammed using the hSTEMCCA polycistronic lentiviral system (*46*). Individual lines were verified for pluripotency through immunofluorescent expression of the markers SSEA4, TRA-1-81, NANOG, and OCT4 (Figure S1). Cells were able to differentiate into the three germ layers by *in vitro* embryoid body (EB) formation (Figure S2), and the microarray-based online PluriTest assay (www.pluritest.org, (*47*)) further confirmed the pluripotency of the lines (Figure S3). Directed neuronal differentiation using the EB intermediate method efficiently produced Nestin-and PAX6-positive neural progenitor cells (NPCs, Figure S4), which further differentiated into βIII-tubulin-positive neurons and GFAP-positive glial cells (Figure S5).

All the patient-derived cells retained the large repeat expansion mutation. Southern blotting revealed bands between 7 and 8.5kb, corresponding to repeat sizes between 750 and 1000 units. Reprogramming to iPS cells and differentiation to NPCs and neurons yielded various fragment sizes suggesting instability of the repeat, both increasing and decreasing in size by hundreds of repeats (Figure S6). Other groups have similarly observed this repeat instability within cell lines, along with somatic mosaicism within affected patients (*4*, *17*, *48*).

### CRISPR design and repeat knockout generation

In order to excise the repeat expansion, we used the CRISPR-Cas9 genome editing system with two guide RNAs designed to target 4 base pairs (bp) upstream and 49 bp downstream of the hexanucleotide repeat (Figure 1a). These sites were selected using a web-based software tool that minimizes potential off-target effects (http://crispr.mit.edu/) (*49*). Each was cloned into the px330 human-optimized CRISPR-Cas9 plasmid (*50*) and introduced to the iPS cells by nucleofection. Subsequent PCR of the region surrounding the repeat showed an amplicon of approximately 65 bp smaller than the normal expected size, indicating simultaneous cleavage and removal of the spanning region by non-homologous end-joining (NHEJ) (Figure 1b).

**Figure 1.**
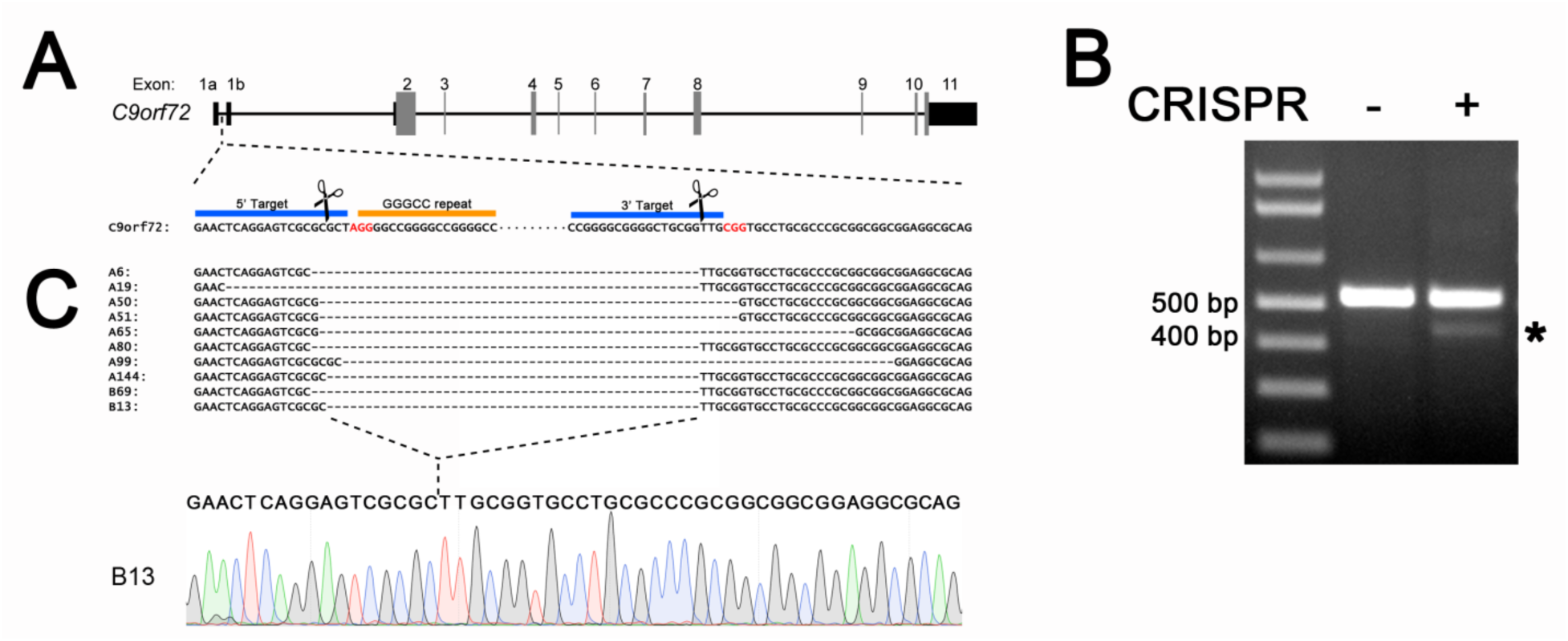
CRISPR-Cas9 design for the deletion of the *C9orf72* repeat expansion mutation. A) Two guide RNAs were used to target sequences (blue bar) flanking the hexanucleotide repeat (orange bar). The Protospacer Adjacent Motif (PAM) is in red. B) PCR of the region surrounding the repeat in iPS cells shows a major amplicon corresponding to the normal allele and a smaller deletion band (asterisk) when cells were treated with both sgRNA-Cas9 constructs. C) Sequence alignment of the deletion band from clones show that the region is targeted, but often contain indels, typical of NHEJ.

Cells were passaged at clonal density and single iPS cell colonies were screened for the presence of the deletion band and the absence of the expansion mutation by Repeat-Primed (RP)-PCR. Out of 593 colonies picked from multiple different parent lines, 66 (11.1%) contained a deletion band. Sequencing of this band showed the repeat region was successfully targeted, though the accuracy of the nuclease and/or repair varied among the clones, with some lines losing as many as 23 additional nucleotides (Figure 1c). Further selection was repeated to obtain homogeneity, until clones were negative for either the wild type amplicon or the stutter pattern in the RP-PCR (Figure 2). 5 lines from 2 patients have been obtained thus far. One of the mutant parent lines (Line A) originally had an expansion of around 845 repeats and a normal allele with 2 repeats (+/exp). From this line, 4 clones were isolated (A50, A51, A65, A80) that were negative for the repeat when tested by RP-PCR (Figure 2b) indicating that the deletion occurred on the expanded allele (+/Δ). Another parent line used in this study had an expansion of 800-1050 units and a normal allele of 5 repeats (Line B). Clone B13 isolated from this parent line was negative for the wild type amplicon and positive in the RP-PCR (Figure 2a-b), indicating a single allelic deletion occurred on the normal allele (Δ/exp). Southern blots confirmed that our CRISPR treatment effectively induced the complete and permanent loss of the *C9orf72* repeat expansion mutation in the 4 clones from parent A (A50, A51, A65, A80), while clone B13 retained the expansion mutation (Figure 2c).

**Figure 2.**
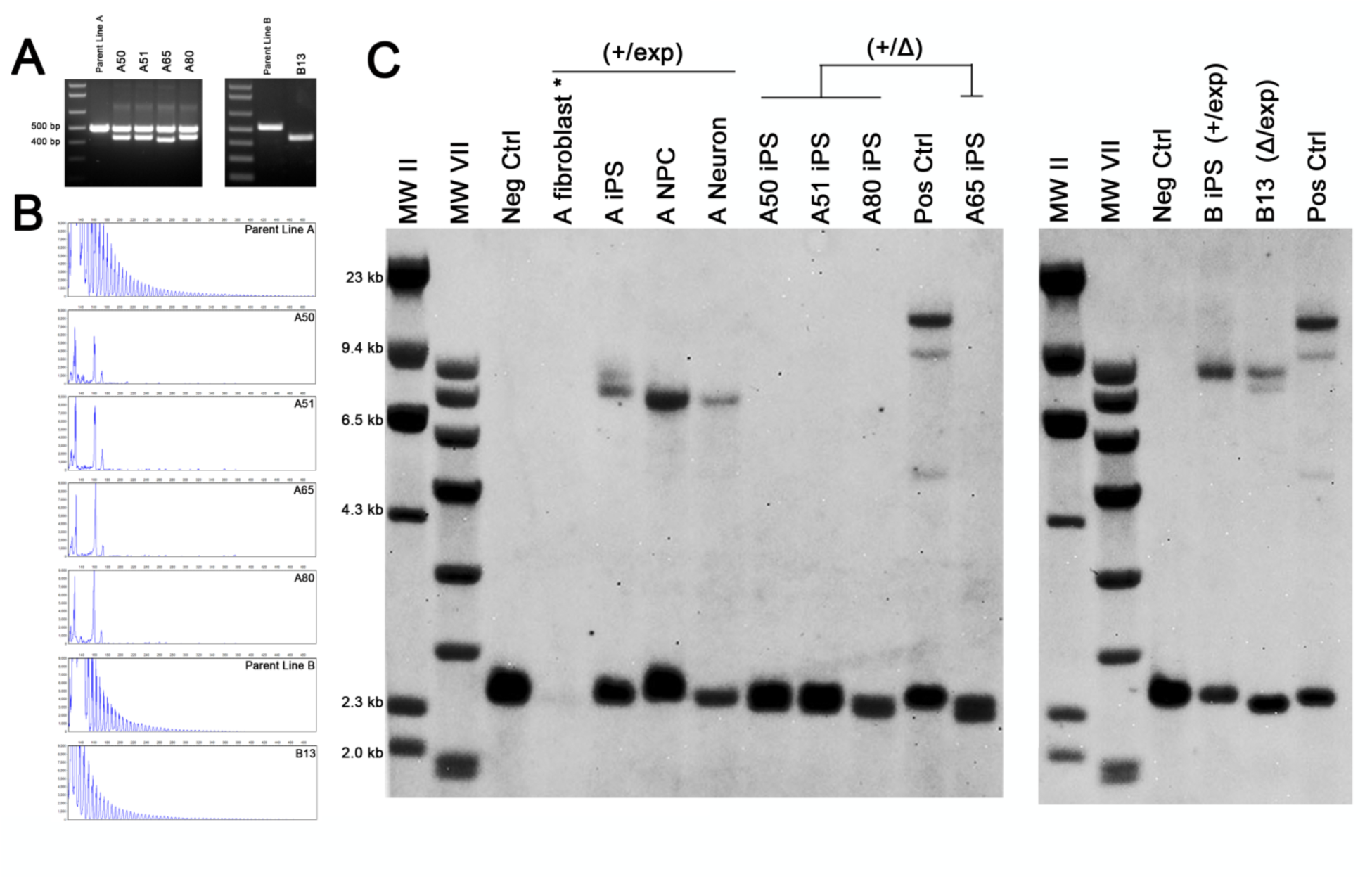
Clonal isolation of repeat knockout iPS cells. A) PCR analysis of four clones (A50, A51, A65, A80) from parent line A containing both a deletion band and an intact normal allele. One clone (B13) from parent line B showed only the deletion band, indicating that the normal allele was targeted. B) RP-PCR showing the loss of the repeat expansion mutation in clones A50, A51, A65, and A80, while the repeat expansion remained in line B13. C) Southern blot confirming that all clones were heterozygous for the deletion. Clones from line A completely lost the repeat expansion mutation (+/Δ) while clone B13 had the deletion occur on the normal allele (Δ/exp). Parent line A fibroblasts (*) were rerun for better visualization in Figure S6.

These repeat knockout (ReKO) iPS cells retained their pluripotency and remained free from karyotype abnormalities (Figure S7). SNP genotyping showed an overall concordance rate of >99.99% (discordance rate of 8.3 e-5) across unedited parent and ReKO lines derived from the same patient. By contrast, the concordance between these and additional genotype data from a number of different patients was on average 74.63%, thus validating that our parent and ReKO lines indeed originated from the same biological specimen.

### No off-target genome editing observed in high-probability cleavage sites

Previous studies have demonstrated that CRISPR-Cas9 RNA-guided nucleases are capable of inducing off-target mutations with a highly variable frequency, sometimes exceeding even those of the intended locus (48). To ensure that the observed phenotypic difference in our ReKO lines are not the epistatic effect of one or more unintended off-target mutations, we performed whole-genome sequencing (WGS) followed by indel calling for comparative analysis between the parent and ReKO-A50 lines. As the specificity of off-target mutagenesis detected by WGS is inherently limited by the baseline accuracy of indel calling of our NGS processing pipeline, we limited our search space to the top 1000 computationally-determined off-target cleavage sites for each guide RNA target sequence, respectively. Only six of the 1,959 top potential off-target sites examined displayed evidence for the putative presence of an indel with respect to the parent line, none of which passed typical quality filters for WGS genotyping.

### Deletion of the repeat expansion corrects RNA foci formation but RAN dipeptides are not observed

One of the cellular hallmarks of the *C9orf72* repeat expansion mutation is the expression of RNA foci through the transcripts initiating upstream of the repeat at exon 1a (*1*, *2*, *17*, *27*, *45*). Consistent with these findings, RNA foci, both in the sense and
antisense direction, were detected in repeat carrier iPS cell lines, as well as differentiated neurons (Figure S8). CRISPR-mediated deletion of the repeat expansion effectively prevented RNA foci formation in these cells, whereas cells with the deletion on the normal unexpanded allele continued to exhibit foci (Figure 3a).

**Figure 3.**
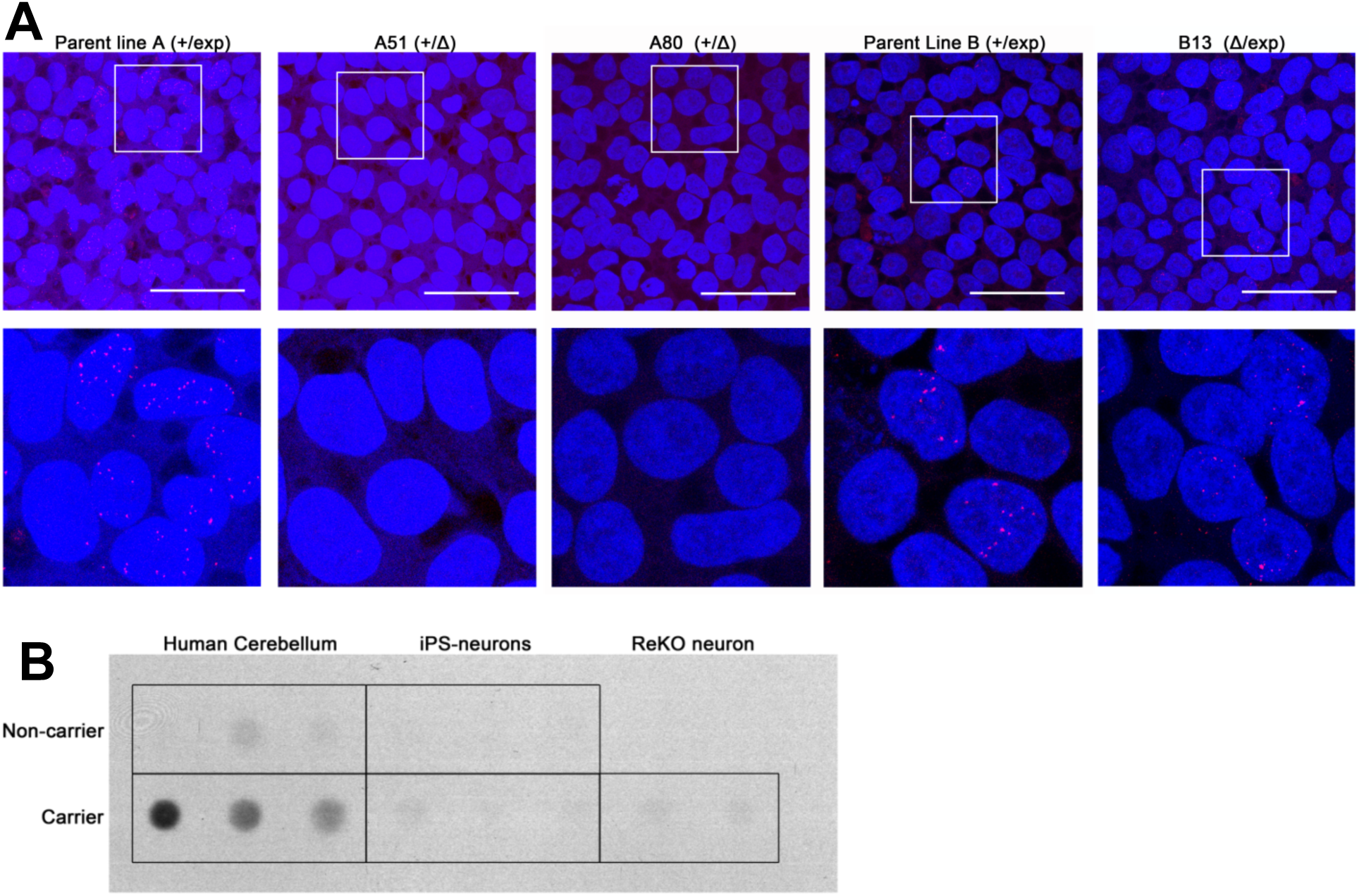
RNA foci and RAN dipeptide expression in *C9orf72* ReKO cells. A) Deletion of the repeat expansion mutation precludes RNA foci formation. Both repeat-carrier (+/exp) parent iPS cell lines exhibit RNA foci formation. Deletion of the expansion mutation (+/Δ) completely prevents expression of foci. Cells with the deletion on the normal allele (Δ/exp) continue to form foci. Scale bar: 50um. B) RAN dipeptide products are not detectable in iPS cell-derived neurons. Dot blot analysis of the GA dipeptide shows detectable levels only in human patient brains with the expansion mutation.

RAN dipeptide deposition is another unique pathological hallmark of *C9orf72*-related neurodegenerative diseases. Aberrant translation of the hexanucleotide repeat generates large repeating dipeptide products that aggregate to form intracellular inclusions in the brains of affected patients (*26*, *28*). In spite of this, RAN dipeptide products were not observed in our iPS cell-derived neurons when examined by immunofluorescence (not shown). Similarly, dot blots were unable to detect any significant RAN dipeptide production in our neuronal cultures (Figure 3b).

### Deletion of the repeat expansion rescues hypermethylation at the C9orf72 locus

Other non-coding repeat expansion mutations are thought to cause disease through epigenetically-mediated downregulation of expression. In the case of Fragile X Syndrome, a CGG repeat expansion in the 5’ untranslated region of *FMR1* leads to DNA hypermethylation and deficiency of the protein (*52*, *53*). Likewise, the *C9orf72* repeat lies within a GC-rich region that harbors CpG islands previously found to be hypermethylated in ˜10-35% of C9-FTD and ALS patient blood samples (*5*, *13*). Bisulfite conversion and sequencing of our cells showed that CpG hypermethylation is indeed present in iPS cells and neurons carrying the expansion mutation (Figure 4). Quantification of the methylation through pyrosequencing showed that the deletion of the expanded repeat resulted in reduction of methylation in all 4 ReKO lines to levels resembling non-carrier controls. Deletion of the normal allele did not reverse the hypermethylation indicating that the effect is due to the expansion of the repeat and is rescued with its deletion by our CRISPR constructs (Figure 4b, Figure S9, Table S1).

**Figure 4.**
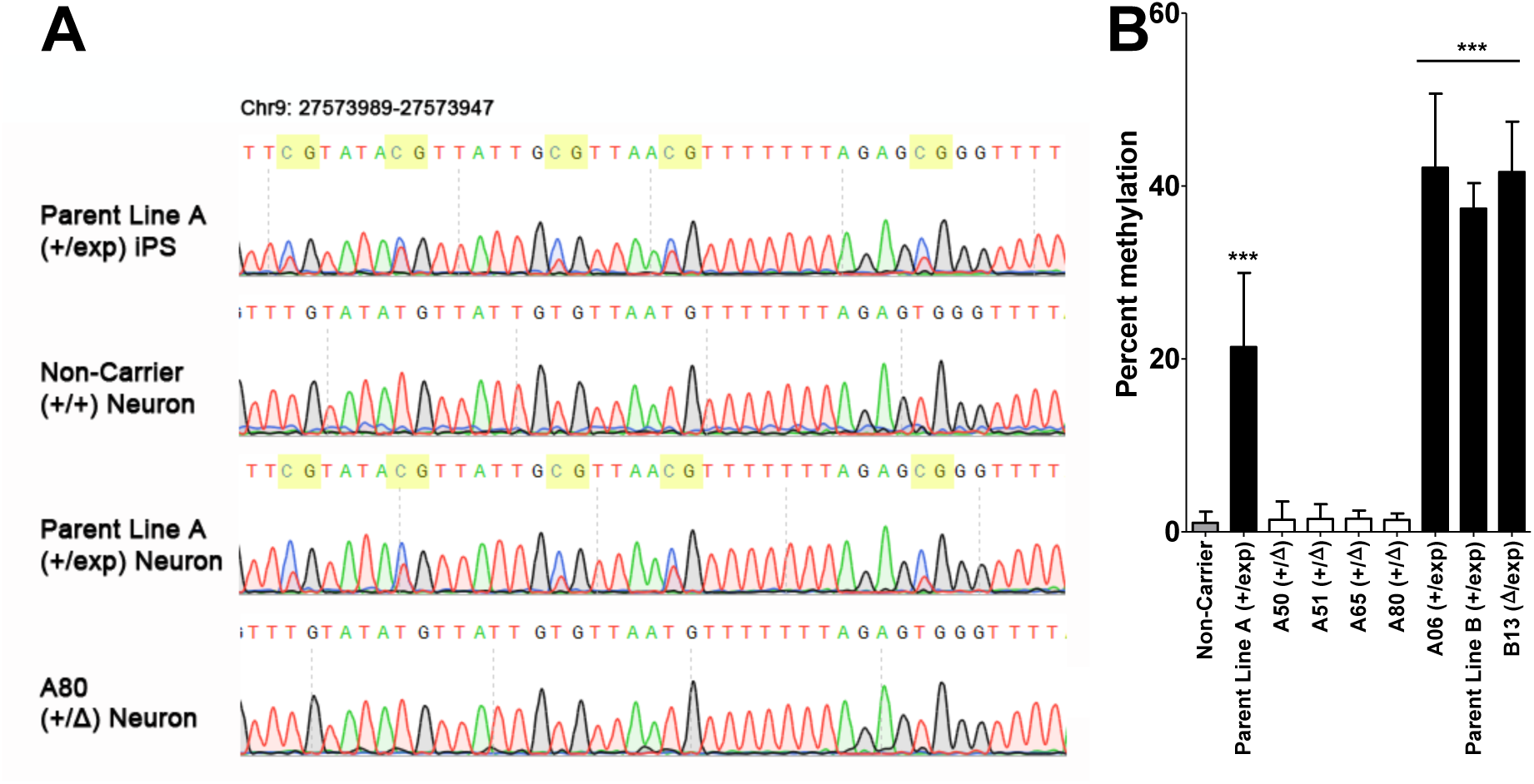
CpG promoter hypermethylation is corrected after deletion of the repeat expansion mutation. A) Bisulfite sequencing of the CpG island upstream of *C9orf72* shows the CpG sites are methylated in repeat carrier iPS cells and derived neurons. B) Pyrosequencing quantification shows overall methylation of the CpG island is increased in repeat expansion carriers. Repeat knockout cells have methylation states restored to non-carrier levels. Clone B13, which has the deletion on the normal allele, and clone A06, which did not have any deletion, continued to have high levels of methylation. Percent methylation is the mean across the 10 CpG sites examined ± S.D.

### Knockout of the C9orf72 repeat expansion mutation does not alter C9orf72 expression

Because our repeat deletion strategy is targeting a regulatory region of *C9orf72*, it is possible that this intervention may disrupt normal expression. If haploinsufficiency of C9orf72 is indeed contributing to disease, further interference with a regulatory region may exacerbate disease progression, especially if the normal allele is also targeted. On the other hand, the repeat expansion itself was reported to be hypermethylated and associated with downregulation of transcription (*12*). If this expansion mutation indeed introduces a negative regulatory element, deletion of this region may be beneficial. C9orf72 protein levels were examined in our iPS cell-derived neurons and no significant difference was observed in neurons containing the repeat mutation (Figure 5a-b). Similarly, transcript levels measured by both qPCR and RNA-seq showed no statistical difference (Figure 5c-d). Others have similarly reported a lack of difference in C9orf72 expression in patient samples and iPS cell-derived neurons (2, *17*). More importantly, CRISPR-mediated deletion of the repeat expansion mutation did not affect transcription or protein expression in each of our independent ReKO neuronal lines, suggesting manipulation of this non-coding region is tolerable.

**Figure 5.**
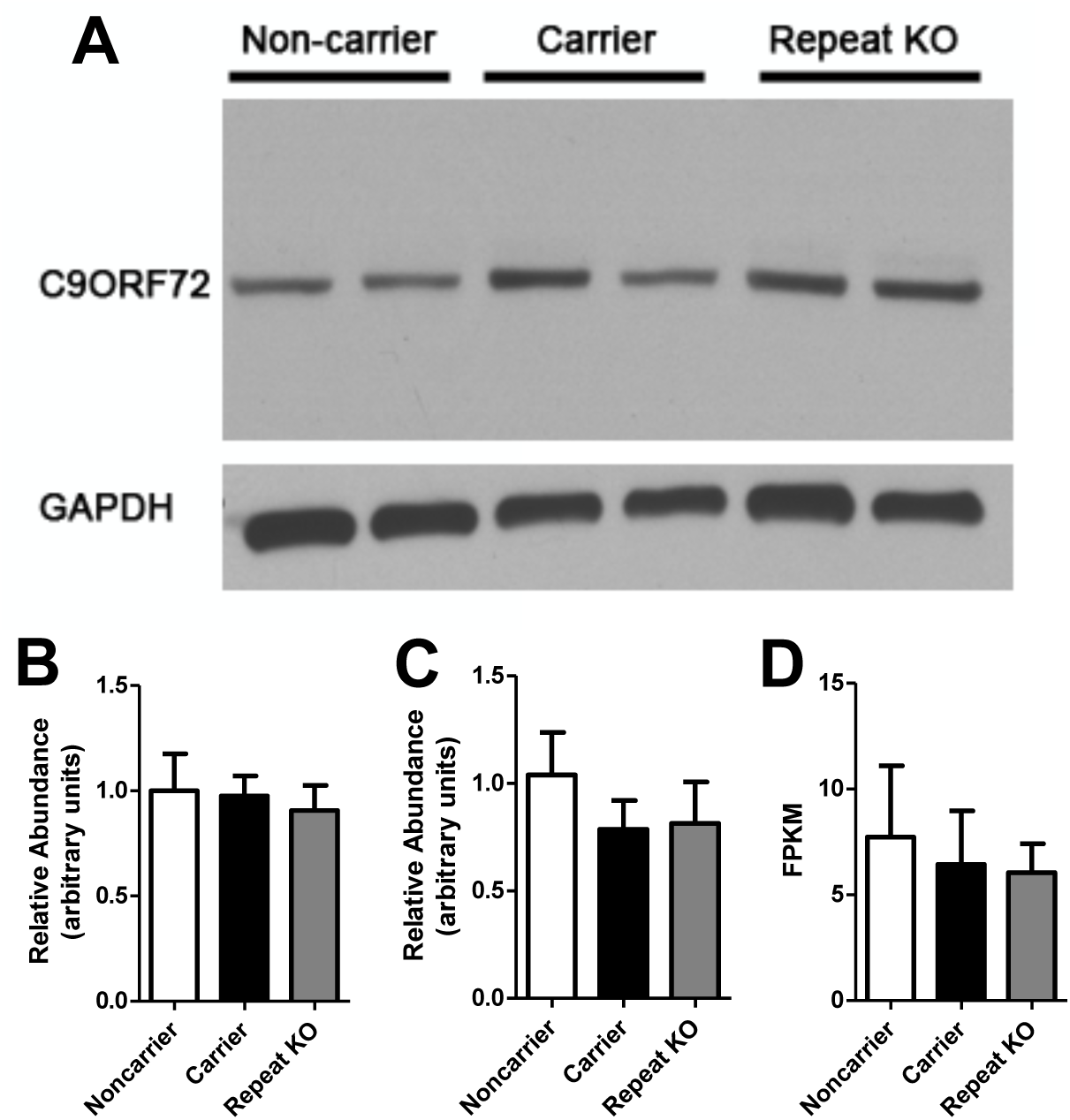
Repeat deletion does not affect C9orf72 expression. A) Representative western blot of iPS cell-derived neurons showing normal expression of C9orf72 among expansion carriers and repeat knockout cells. B) Quantification of western blots normalized to GAPDH and analyzed by one-way ANOVA (mean ± S.D., n=3 per genotype). C) Taqman qPCR assay (Hs00376619_m1) of total *C9orf72* transcription show no significant differences by one-way ANOVA (mean ± S.D., n=3 per genotype). D) FPKM levels of C9orf72 from RNAseq of noncarrier (n=13), carrier (n=14), and repeat KO (n=3) neurons show no significant difference. One-way ANOVA was performed to determine significance (F = 0.94, p = 0.43).

## Discussion

Here we demonstrated the capability of the CRISPR-Cas9 system to permanently delete the large repeat expansion mutation in *C9orf72*-associated disease. Using two guide RNAs targeting regions flanking the repeat expansion, simultaneous cleavage was able to remove a full expansion mutation, completely prevent formation of RNA foci, and rescue local hypermethylation. Multiple lines were generated targeting both alleles, but none of the clones isolated thus far contained homozygous deletions. The deleted allele often contained small additional indels typical of repair by NHEJ (*54*), but because the targeted region is within an intronic region away from splice sites, these deletions may have little or no effect on gene expression. Indeed, loss of this repeat region did not significantly alter expression of C9orf72, which may be of relevance given the recent discovery of its role in microglia and inflammation (*55*). The rescue from promoter hypermethylation is particularly interesting because multiple studies have reported its function in reducing expression of *C9orf72* (*5*, *12*, *13*). Although this downregulation is hypothesized to cause disease through haploinsufficiency, it has also been suggested to be neuroprotective through the consequential reduction in RNA foci and RAN dipeptide production (*56*, *57*). If in the diseased patient this hypermethylation does contribute to pathogenesis, then targeting the repeat expansion for deletion using CRISPRs may be a more favorable strategy. Attempts to target the repeat using antisense oligonucleotides would not be able to rescue this haploinsufficiency (*17*), and simultaneously targeting both the sense and antisense products of the repeat may be difficult given their complementary sequence. Our results here show that RNA foci expression and promoter hypermethylation can be blocked while preserving endogenous protein expression.

Although our CRISPR strategy was able to preserve *C9orf72* expression, we did not observe an increase in expression with the rescue of promoter hypermethylation as expected. Actually, we did not observe any deficit in transcript or protein expression in our *C9orf72* mutant cell lines despite having hypermethylation. This discrepancy remains unclear, but it does suggest a more complex mechanism of transcriptional regulation than methylation alone. Together with the lack of RAN dipeptide expression, this suggests the mutation alone is not sufficient to induce haploinsufficiency or RAN dipeptide deposition, at least in our cell culture system, though it is important to note that iPS cell-derived neurons resemble fetal neurons and may not be the best model for the aged diseased brain (*58*). It is possible the effects of the mutation may take decades to accumulate or may arise in a non-cell-autonomous manner. Our cells did however exhibit the RNA foci that are characteristic of these repeat expansion disorders, but their presence in these viable embryonic-like cells argues against a primary role in pathogenesis. Nonetheless, they do provide a valuable readout for the presence and transcription of the repeat expansion mutation.

Regardless of whether the *C9orf72* repeat expansion mutation acts through a gain-of-toxicity or loss-of-function, our strategy of deleting the entire repeat expansion itself may resolve both. Furthermore, if disease severity is indeed correlated with repeat size, removing the entire repeat should prevent the unstable somatic expansion that is common in these repeat diseases. We demonstrated the feasibility of targeted genomic editing to excise the repeat expansion region, reversing cellular phenotypes while preserving *C9orf72* mRNA and protein expression *in vitro*. Whether this mutation is a viable target *in vivo* is a question for future studies when models of disease become available. BAC mice exhibit RNA foci and RAN dipeptide deposition but not behavioral
deficits or neurodegeneration (*39*, *40*). Targeting the repeat *in vivo* would likely require Adeno-Associated Viral (AAV) delivery, which has low immunogenicity, has already been demonstrated in murine CNS (*59*), and is already used in the clinic (*60*, *61*). This strategy of deleting repeat expansions may be applicable to other similar diseases with intronic non-coding repeats, such as Fragile X or Friedreich’s Ataxia. The use of NHEJ, though more efficient than homologous recombination, often produces indels, which may be disastrous if not for its occurrence in non-coding regions, and we have shown that the normal gene expression can be preserved. Translation into the clinic would still follow the advancement of CRISPR technology, when improvements in genome editing and safety are demonstrated in patients.

## Materials and Methods

### Induced pluripotent stem cell generation

Fibroblasts from healthy controls and FTD patients with the *C9orf72* repeat expansion were obtained from the UCSF Memory and Aging Center with patient consent and IRB approval (#10-000574). Reprogramming was performed under UCLA ESCRO approval (#2013-001-01) using the hSTEMCCA polycistonic lentiviral vector (Millipore) (*46*), and maintained on Matrigel (BD Biosciences) in mTeSR media (Stem Cell Technologies). Pluripotency was confirmed by immunofluorescence of markers, in vitro embryoid body (EB) formation, and the microarray-based Pluritest assay (*47*). For EB formation, colonies were dissociated using dispase (Life Technologies) and plated onto ultra-low attachment plates (Corning) in media lacking FGF. Differentiated EBs were embedded in Histogel (Fisher) prior to
paraffin embedding and sectioning. For the Pluritest assay, RNA was extracted from iPS cells free from any spontaneous differentiation and run on an Illumina Human HT-12 v4 Expression array by the UCLA Neuroscience Genomics Core. Raw data files were uploaded to the Pluritest website (www.pluritest.org) for analysis. Karyotyping was performed by Cell Line Genetics and monitored around every 25 passages.

### Antibodies

The following antibodies were used: Nanog (Cell Signaling, D73G4, 1:500), SSEA4 (BD Biosciences, MC813-70, 1:250), OCT3/4 (Santa Cruz Biotechnology, H-134, 1:250), TRA-1-81(Stem Cell Technologies, 60065, 1:500), Alpha-fetoprotein (Thermo Fisher, SP154, 1:100), Nestin (Thermo Fisher, 10C2, (1:250), Smooth Muscle Actin (Thermo Fisher, 4A4, 1:500), GFAP (Antagene, V1058, 1:1000), βIII-Tubulin (Covance, PRB-435P, 1:2500) C9ORF72 (Novus, NBP1-93504, 1:500), poly-GA (Cosmo Bio, TIP-C9-P01, 1:1000), GAPDH (Santa Cruz Biotechnology, sc-365062, 1:1000), ZO1 (Thermo Fisher, 61-7300, 1:250).

### CRISPR design

The px330 plasmid (#42230, Addgene), which contains the Streptococcus pyogenes Cas9 (SpCas9) nuclease along with the guide RNA, was used for CRISPR-Cas9 deletion of the repeat. For the upstream target, oligos [5’-caccGAACTCAGGAGTCGCGCGCT-3’] and [5’-aaacAGCGCGCGACTCCTGAGTTC-3’]were annealed and cloned into the BbsI-digested plasmid. For the downstream target,oligos [5’-caccgCGGGGCGGGGCTGCGGTTG-3’] and [5’-aaacCAACCGCAGCCCCGCCCCGC-3’] were similarly cloned into px330. Cells were dissociated using Accutase (Life Technologies) and 2 ug of each plasmid was used to nucleofect 1x106 cells using the Amaxa Human Stem Cell Nucleofector^®^ Kit 2 (Lonza) with program A-23. Cells were plated at clonal density using ReLeSR (Stem Cell Technologies) and Y-27632 (10 uM, Stemgent). Colonies were screened for the deletion by PCR using the primers: fwd 5’-GAGGAGAGCCCCCGCTTCTAC-3’, rev 5’-CCGCTAGGAAAGAGAGGTGCG-3’. Further clonal selection was performed as necessary.

### Repeat expansion genotyping

The presence of the *C9orf72* repeat expansion was determined using the repeat-primed PCR assay as described in (1). Southern blotting was performed as described in (1). The size standards used for Southern blots are DNA MW Marker II (23130, 9416, 6557, 4361, 2322, and 2027 bp), and DNA MW Marker VII (8576, 7427, 6106, 4899, 3639, 2799, 1953, 1882, 1515, 1482, 1164 bp) (both Roche).

### Neuronal differentiation

Neuronal differentiation was performed using the StemDIFF Neural system (Stem Cell Technologies). 2x10^6^ iPS cells were dissociated with Accutase and plated onto Agrewell 800 plates with Neural Induction Media (Stem Cell Technologies) and ROCK inhibitor (10 uM Y-27632. Stemgent). After 5 days, embryoid bodies were plated onto poly-ornithine/laminin-coated plates to allow for rosette formation which were isolated using the Rosette Selection Media (Stem Cell Technologies) on day 7 following plating. The neural progenitor cells were passaged using Accutase and maintained on Matrigel-coated plates in N2/B27 Media (1x DMEM/F12 basal media, 0.5x N2, 0.5x B27, all Life Technologies) supplemented with EGF and FGF (both 10ng/mL, Stemgent). Further differentiation to neurons was performed by passaging onto poly-ornithine/laminin-coated plates in N2/B27 Media supplemented with 1uM cAMP (Sigma), 200 ng/mL Ascorbic Acid (Sigma), and 10 ng/mL BDNF (Peprotech). Neurons were allowed to mature for at least 4 weeks.

### Quantitative RT-PCR

RNA was harvested with the RNEasy Plus Mini Kit (Qiagen) and converted to cDNA using Superscript III First Strand Synthesis System (Life Technologies). QPCR of total *C9orf72* was performed using Taqman (assay Hs00376619_m1) and amplification efficiencies and Cp values were acquired using a Roche 480 LightCycler. Relative abundance was calculated using the Pfaffl method against three reference genes, *GAPDH* (Hs02758991_g1), *ACTB* (Hs01060665_g1), and *RPLP0* (Hs99999902_m1).

### RNAseq

RNA sequencing was performed by the UCLA Neuroscience Genomics Core. Libraries were prepared using the Illumina TruSeq Stranded Total RNA with Ribo-Zero Gold kit. Paired-end 69 bp reads were acquired using an Illumina Hiseq2500, with four samples per lane per run, corresponding to ˜40 million reads per sample. FPKM was estimated from count-level data. One-way ANOVA was performed using oneway.test in R v3.2.0.

### Protein extraction and blotting

Cells were lysed in RIPA buffer (Cell Signaling Technology) and cleared by centrifugation at 20000g. Frozen human autopsy cerebellar samples from C9-positive FTD patients and controls were homogenized similarly with a dounce homogenizer. For dot blot assays, cell pellets were homogenized using 8M urea. Protein lysates were quantified by BCA assay (Biorad) and spotted on a dot blot or separated by electrophoresis prior to western transfer. Quantification was done manually using ImageJ.

### Fluorescence in situ hybridization (FISH)

FISH was performed as described in (17). Briefly, cells on coverslips were fixed in 4% paraformaldehyde (Electron Microscopy Services) and permeabilized using 0.25% Triton X-100. Cells were washed using RNAse-Free PBS and pre-hybidized with hybridization buffer (50% formamide, 2X SSC, 10% Dextran Sulfate, 50 mM sodium phosphate pH 7.0) for 30 min at 66°C. 50nM of the FISH probe was hybridized for 3 hours at 66°C in the dark. The probe for detecting sense foci is CCCCGG4-Quasar570 (Biosearch Technologies) or TYE563-CCCCGGCCCCGGCCCC (Exiqon). The probe for detecting antisense foci is TYE563-GGGGCCGGGGCCGGGG (Exiqon). Cells were washed once with 2X SSC/0.1% Tween-20 at room temperature followed by three stringent washes in 0.2x SSC at 66°C. For co-immunofluorescence experiments, cells were immediately incubated with primary antibodies in RNAse-free PBST for 1 hour at room temperature followed by the secondary for an additional hour. Nuclei were counterstained using TO-PRO-3 (Life Technologies) and mounted in ProLong Gold (Life Technologies) or in Vectashield with DAPI (Vector Labs). Image stacks were acquired using a Zeiss LSM 5 Pascal confocal microscope and processed in ImageJ. Foci were manually counted.

### Methylation assays

Genomic DNA was bisulfite converted using the EpiTect Fast Bisulfite Conversion Kit (Qiagen) as instructed. The CpG islands upstream of the repeat expansion were amplified by PCR with the KAPA HiFi Uracil+ Polymerase (Kapa Biosystems). Primers and cycling parameters were done as described in Xi *et al*. 2013 (*57*). Products were sequenced and methylated CpGs were defined as having (T/C) double peaks. For quantification of methylation, genomic DNA was sent to EpigenDX for bisulfite conversion and pyrosequencing of the CpG island upstream of Exon 1a (Assay ADS3233-FS1 and ADS3232-FS1). One-way ANOVA with Bonferroni’s correction was performed in Prism (Graphpad).

### SNP Genotyping and Whole Genome Sequencing

Genotype data was generated using the Infinium Omni2.5Exome-8 Beadchip (Illumina) using genomic DNA extracted from two parent and two ReKO lines derived from the same patient, as well as from whole-blood draws from four randomly selected individuals, according to manufacturer’s specifications. Concordance on all autosomal SNP called across all samples (n=2,512,633) was calculated using SNP & Variation Suite software (Golden Helix).

Whole genome sequencing for the parent and ReKO lines was performed by the New York Genome Center. Library preparation was performed using the Illumina TruSeqDNA PCR-free protocol, to generate 150 bp paired-end reads. Sequencing was performed using the Illumina HiSeqX to an average depth of 30x autosomal coverage. Read alignment was performed to the NCBI GRCh37 reference genome using BWA. Duplicate reads were removed by Picard. Base quality score recalibration, indel realignment, and downstream variant calling was performed using GATK.

Indel calls were left-aligned to account for genomic ambiguity and VCF tools was used to distinguish indel calls only present in the ReKO A50 sample. We used CROP-IT (http://cheetah.bioch.virginia.edu/AdliLab/CROP-IT/homepage.html) (58) for in-silico determination of the top 1000 potential off-target cleavage sites for each guide RNA. To account for the heterogeneity (59), each putative site was expanded by 100 bp on each side, and overlapping regions were merged. This resulted in 1959 separate loci which were screened for indel calls unique to CRISPR-Cas9 treated ReKO A50 line.

**Supplementary Materials**

**Supplementary Figures**

**Figure S1**. Normal iPS cell generation from *C9orf72* expansion carriers.

**Figure S2**. iPS cells are pluripotent.

**Figure S3**. All iPS cells pass the Pluritest assay.

**Figure S4**. Efficient generation of neural progenitor cells from iPS generated from C9orf72 expansion carriers.

**Figure S5**. Directed neuronal differentiation.

**Figure S6**. Southern blot of iPS and differentiated cells shows instability in repeat size.

**Figure S7**. Pluripotency is maintained with CRISPR-mediated deletion of the *C9orf72* repeat.

**Figure S8**. Repeat Expansion carrier cells express RNA foci.

**Figure S9**. Pyrosequencing quantification of CpG island methylation.

**Supplementary Table**

**Table S1**. Quantification of *C9orf72* promoter CpG sites by pyrosequencing.

## Acknowledgments

We would like to thank William X Yang, and members of the Coppola laboratory for critical reading of the manuscript and helpful comments, and the UCLA Neuroscience Genomics Core for generating microarray and RNA-sequencing data.

### Funding

This work was supported by the NIH (NIH/National Center for Advancing Translational Science (NCATS) UCLA CTSI Grant Number UL1TR000124, RC1 AG035610), the Tau Consortium, and the John Douglas French Alzheimer’s Foundation. We acknowledge the support of the NINDS Informatics Center for Neurogenetics and Neurogenomics (P30 NS062691).

### Author contributions

MP, ZY, and GC designed the study; MP, ZY, TSK, EWS, DD, JB, ATF, KW, QW, EL, SN, and FFC performed experiments and generated data; AK, JF, HVV, and BLM enrolled patients and collected biological samples; AYH, JAC, FG, performed statistical analyses; MP and GC wrote the manuscript; MP, EWS, JAC, HVV, and GC revised the manuscript.

### Competing interests

JAC is a co-founder and has an equity interest in Verge Genomics, a genomics-based biotechnology company.

## References and Notes

1. M. DeJesus-Hernandez et al., Expanded GGGGCC hexanucleotide repeat in noncoding region of C9ORF72 causes chromosome 9p-linked FTD and ALS. Neuron 72, 245 (Oct 20, 2011).

2. A. E. Renton et al., A hexanucleotide repeat expansion in C9ORF72 is the cause of chromosome 9p21-linked ALS-FTD. Neuron 72, 257 (Oct 20, 2011).

3. T. P. Levine, R. D. Daniels, A. T. Gatta, L. H. Wong, M. J. Hayes, The product of C9orf72, a gene strongly implicated in neurodegeneration, is structurally related to DENN Rab-GEFs. Bioinformatics 29, 499 (Feb 15, 2013).

4. M. van Blitterswijk et al., Association between repeat sizes and clinical and pathological characteristics in carriers of C9ORF72 repeat expansions (Xpansize-72): a cross-sectional cohort study. Lancet neurology 12, 978 (Oct, 2013).

5. I. Gijselinck et al., The C9orf72 repeat size correlates with onset age of disease, DNA methylation and transcriptional downregulation of the promoter. Molecular psychiatry, (Oct 20, 2015).

6. I. Gijselinck et al., A C9orf72 promoter repeat expansion in a Flanders-Belgian cohort with disorders of the frontotemporal lobar degeneration-amyotrophic lateral sclerosis spectrum: a gene identification study. Lancet neurology 11, 54 (Jan, 2012).

7. A. J. Waite et al., Reduced C9orf72 protein levels in frontal cortex of amyotrophic lateral sclerosis and frontotemporal degeneration brain with the C9ORF72 hexanucleotide repeat expansion. Neurobiology of aging 35, 1779 e5 (Jul, 2014).

8. S. Xiao et al., Isoform-specific antibodies reveal distinct subcellular localizations of C9orf72 in amyotrophic lateral sclerosis. Annals of neurology 78, 568 (Oct, 2015).

9. C. J. Donnelly et al., RNA toxicity from the ALS/FTD C9ORF72 expansion is mitigated by antisense intervention. Neuron 80, 415 (Oct 16, 2013).

10. M. van Blitterswijk et al., Novel clinical associations with specific C9ORF72 transcripts in patients with repeat expansions in C9ORF72. Acta neuropathologica, (Oct 5, 2015).

11. S. Ciura et al., Loss of function of C9orf72 causes motor deficits in a zebrafish model of amyotrophic lateral sclerosis. Annals of neurology 74, 180 (Aug, 2013).

12. Z. Xi et al., The C9orf72 repeat expansion itself is methylated in ALS and FTLD patients. Acta neuropathologica 129, 715 (May, 2015).

13. Z. Xi et al., Hypermethylation of the CpG island near the G4C2 repeat in ALS with a C9orf72 expansion. American journal of human genetics 92, 981 (Jun 6, 2013).

14. V. V. Belzil et al., Reduced C9orf72 gene expression in c9FTD/ALS is caused by histone trimethylation, an epigenetic event detectable in blood. Acta neuropathologica 126, 895 (Dec, 2013).

15. M. Therrien, G. A. Rouleau, P. A. Dion, J. A. Parker, Deletion of C9ORF72 results in motor neuron degeneration and stress sensitivity in C. elegans. PloS one 8, e83450 (2013).

16. M. Koppers et al., C9orf72 ablation in mice does not cause motor neuron degeneration or motor deficits. Annals of neurology 78, 426 (Sep, 2015).

17. D. Sareen et al., Targeting RNA foci in iPSC-derived motor neurons from ALS patients with a C9ORF72 repeat expansion. Science translational medicine 5, 208ra149 (Oct 23, 2013).

18. M. B. Harms et al., Lack of C9ORF72 coding mutations supports a gain of function for repeat expansions in amyotrophic lateral sclerosis. Neurobiology of aging 34, 2234 e13 (Sep, 2013).

19. R. N. Kanadia et al., A muscleblind knockout model for myotonic dystrophy. Science 302, 1978 (Dec 12, 2003).

20. Y. B. Lee et al., Hexanucleotide repeats in ALS/FTD form length-dependent RNA foci, sequester RNA binding proteins, and are neurotoxic. Cell reports 5, 1178 (Dec 12, 2013).

21. J. Cooper-Knock et al., Sequestration of multiple RNA recognition motif-containing proteins by C9orf72 repeat expansions. Brain: a journal of neurology 137, 2040 (Jul, 2014).

22. K. Zhang et al., The C9orf72 repeat expansion disrupts nucleocytoplasmic transport. Nature 525, 56 (Sep 3, 2015).

23. A. R. Haeusler et al., C9orf72 nucleotide repeat structures initiate molecular cascades of disease. Nature 507, 195 (Mar 13, 2014).

24. H. Tran et al., Differential Toxicity of Nuclear RNA Foci versus Dipeptide Repeat Proteins in a Drosophila Model of C9ORF72 FTD/ALS. Neuron 87, 1207 (Sep 23, 2015).

25. J. D. Cleary, L. P. Ranum, Repeat-associated non-ATG (RAN) translation in neurological disease. Human molecular genetics 22, R45 (Oct 15, 2013).

26. P. E. Ash et al., Unconventional translation of C9ORF72 GGGGCC expansion generates insoluble polypeptides specific to c9FTD/ALS. Neuron 77, 639 (Feb 20, 2013).

27. T. F. Gendron et al., Antisense transcripts of the expanded C9ORF72 hexanucleotide repeat form nuclear RNA foci and undergo repeat-associated non-ATG translation in c9FTD/ALS. Acta neuropathologica, (Oct 16, 2013).

28. K. Mori et al., The C9orf72 GGGGCC repeat is translated into aggregating dipeptide-repeat proteins in FTLD/ALS. Science 339, 1335 (Mar 15, 2013).

29. Y. J. Zhang et al., Aggregation-prone c9FTD/ALS poly(GA) RAN-translated proteins cause neurotoxicity by inducing ER stress. Acta neuropathologica 128, 505 (Oct, 2014).

30. S. May et al., C9orf72 FTLD/ALS-associated Gly-Ala dipeptide repeat proteins cause neuronal toxicity and Unc119 sequestration. Acta neuropathologica 128, 485 (Oct, 2014).

31. S. Mizielinska et al., C9orf72 repeat expansions cause neurodegeneration in Drosophila through arginine-rich proteins. Science 345, 1192 (Sep 5, 2014).

32. X. Wen et al., Antisense proline-arginine RAN dipeptides linked to C9ORF72-ALS/FTD form toxic nuclear aggregates that initiate in vitro and in vivo neuronal death. Neuron 84, 1213 (Dec 17, 2014).

33. D. Yang et al., FTD/ALS-associated poly(GR) protein impairs the Notch pathway and is recruited by poly(GA) into cytoplasmic inclusions. Acta neuropathologica 130, 525 (Oct, 2015).

34. I. R. Mackenzie et al., Quantitative analysis and clinico-pathological correlations of different dipeptide repeat protein pathologies in C9ORF72 mutation carriers. Acta neuropathologica, (Sep 15, 2015).

35. I. R. Mackenzie et al., Dipeptide repeat protein pathology in C9ORF72 mutation cases: clinico-pathological correlations. Acta neuropathologica 126, 859 (Dec, 2013).

36. Y. S. Davidson et al., Brain distribution of dipeptide repeat proteins in frontotemporal lobar degeneration and motor neurone disease associated with expansions in C9ORF72. Acta neuropathologica communications 2, 70 (2014).

37. P. Gami et al., A 30-unit hexanucleotide repeat expansion in C9orf72 induces pathological lesions with dipeptide-repeat proteins and RNA foci, but not TDP-43 inclusions and clinical disease. Acta neuropathologica 130, 599 (Oct, 2015).

38. M. Proudfoot et al., Early dipeptide repeat pathology in a frontotemporal dementia kindred with C9ORF72 mutation and intellectual disability. Acta neuropathologica 127, 451 (Mar, 2014).

39. O. M. Peters et al., Human C9ORF72 Hexanucleotide Expansion Reproduces RNA Foci and Dipeptide Repeat Proteins but Not Neurodegeneration in BAC Transgenic Mice. Neuron 88, 902 (Dec 2, 2015).

40. J. G. O’Rourke et al., C9orf72 BAC Transgenic Mice Display Typical Pathologic Features of ALS/FTD. Neuron 88, 892 (Dec 2, 2015).

41. E. Majounie et al., Frequency of the C9orf72 hexanucleotide repeat expansion in patients with amyotrophic lateral sclerosis and frontotemporal dementia: a cross-sectional study. Lancet neurology 11, 323 (Apr, 2012).

42. M. Jinek et al., A programmable dual-RNA-guided DNA endonuclease in adaptive bacterial immunity. Science 337, 816 (Aug 17, 2012).

43. C. E. Nelson et al., In vivo genome editing improves muscle function in a mouse model of Duchenne muscular dystrophy. Science 351, 403 (Jan 22, 2016).

44. C. Long et al., Postnatal genome editing partially restores dystrophin expression in a mouse model of muscular dystrophy. Science 351, 400 (Jan 22, 2016).

45. M. Tabebordbar et al., In vivo gene editing in dystrophic mouse muscle and muscle stem cells. Science 351, 407 (Jan 22, 2016).

46. C. A. Sommer et al., Induced pluripotent stem cell generation using a single lentiviral stem cell cassette. Stem Cells 27, 543 (Mar, 2009).

47. F. J. Muller et al., A bioinformatic assay for pluripotency in human cells. Nature methods 8, 315 (Apr, 2011).

48. S. Almeida et al., Modeling key pathological features of frontotemporal dementia with C9ORF72 repeat expansion in iPSC-derived human neurons. Acta neuropathologica 126, 385 (Sep, 2013).

49. P. D. Hsu et al., DNA targeting specificity of RNA-guided Cas9 nucleases. Nature biotechnology 31, 827 (Sep, 2013).

50. L. Cong et al., Multiplex genome engineering using CRISPR/Cas systems. Science 339, 819 (Feb 15, 2013).

51. Y. Fu et al., High-frequency off-target mutagenesis induced by CRISPR-Cas nucleases in human cells. Nature biotechnology 31, 822 (Sep, 2013).

52. M. V. Bell et al., Physical mapping across the fragile X: hypermethylation and clinical expression of the fragile X syndrome. Cell 64, 861 (Feb 22, 1991).

53. M. Pieretti et al., Absence of expression of the FMR-1 gene in fragile X syndrome. Cell 66, 817 (Aug 23, 1991).

54. W. T. Hendriks, C. R. Warren, C. A. Cowan, Genome Editing in Human Pluripotent Stem Cells: Approaches, Pitfalls, and Solutions. Cell stem cell 18, 53 (Jan 7, 2016).

55. J. G. O’Rourke et al., C9orf72 is required for proper macrophage and microglial function in mice. Science 351, 1324 (Mar 18, 2016).

56. C. T. McMillan et al., C9orf72 promoter hypermethylation is neuroprotective: Neuroimaging and neuropathologic evidence. Neurology 84, 1622 (Apr 21, 2015).

57. E. Y. Liu et al., C9orf72 hypermethylation protects against repeat expansion-associated pathology in ALS/FTD. Acta neuropathologica 128, 525 (Oct, 2014).

58. S. Mahmoudi, A. Brunet, Aging and reprogramming: a two-way street. Current opinion in cell biology 24, 744 (Dec, 2012).

59. F. A. Ran et al., In vivo genome editing using Staphylococcus aureus Cas9. Nature 520, 186 (Apr 9, 2015).

60. L. Lisowski, S. S. Tay, I. E. Alexander, Adeno-associated virus serotypes for gene therapeutics. Current opinion in pharmacology 24, 59 (Oct, 2015).

61. T. Gaj, B. E. Epstein, D. V. Schaffer, Genome Engineering Using Adeno-associated Virus: Basic and Clinical Research Applications. Molecular therapy: the journal of the American Society of Gene Therapy, (Sep 16, 2015).

62. Z. Xi et al., Hypermethylation of the CpG Island Near the GC Repeat in ALS with a C9orf72 Expansion. American journal of human genetics, (May 22, 2013).

63. R. Singh, C. Kuscu, A. Quinlan, Y. Qi, M. Adli, Cas9-chromatin binding information enables more accurate CRISPR off-target prediction. Nucleic acids research 43, e118 (Oct 15, 2015).

64. C. Kuscu, S. Arslan, R. Singh, J. Thorpe, M. Adli, Genome-wide analysis reveals characteristics of off-target sites bound by the Cas9 endonuclease. Nature biotechnology 32, 677 (Jul, 2014).

